# DEGoldS: a workflow to assess the accuracy of differential expression analysis pipelines through gold-standard construction

**DOI:** 10.1101/2022.09.13.507753

**Authors:** Mikel Hurtado, Fernando Mora-Márquez, Álvaro Soto, Daniel Marino, Pablo G. Goicoechea, Unai López de Heredia

**Affiliations:** Departamento de Ciencias Forestales. NEIKER-BRTA, Instituto Vasco de Investigación y Desarrollo Agrario, Campus Agroalimentario de Arkaute, Crtra N-104 km 355, 01192 Arkaute, Alava, España; Dpto. Sistemas y Recursos Naturales. ETSI Montes, Forestal y del Medio Natural. Universidad Politécnica de Madrid. Ciudad Universitaria s/n, 28040 Madrid, Spain; Departamento de Biología Vegetal y Ecología. Facultad de Ciencia y Tecnología. Universidad del País Vasco-Euskal Herriko Unibertsitatea (UPV-EHU). Barrio Sarriena s/n 48940 Leioa, Bizkaia; IKERBASQUE, Basque Foundation for Science, Plaza Euskadi 5,48009 Bilbao, Spain

**Keywords:** RNA-seq, transcriptome, benchmarking, non-model species, assembly

## Abstract

RNA sequencing (RNA-seq) is a high throughput sequencing method that has become one the most employed tools in transcriptomics. The implementation of optimal bioinformatic analyses required in RNA-seq experiments may be complicated due to the large amounts of data generated by the sequencing platforms, along with the intrinsic nature of these data types. In the last years many programs and pipelines have been developed for differential expression (DE) analyses, but their effectiveness can be reduced when working with non-model species lacking public genomic resources. Moreover, there is not a universal recipe for all the experiments and datasets and the modification of standard RNA-seq bioinformatic pipelines through parameter tuning and the use of alternative software may have a strong impact in the outcome of DE analysis. Therefore, although the selection of the most accurate DE pipeline configuration and the evaluation of how these changes could affect the final DE results in RNA-seq experiments is mandatory to reduce bias, the lack of gold-standard datasets with known expression patterns hampers its implementation. In the present manuscript we present DEGoldS, a workflow consisting on sequential Bash and R scripts to construct gold-standards for simulation-based benchmarking of user selected pipelines for DE analysis and the computation of the accuracy of the pipelines. We validated the workflow with a case study consisting on real RNA-seq libraries of radiata pine, an important forest tree species with no publicly available reference genome. The results showed that slight pipeline modifications produced remarkable differences in the outcome of DE analysis.

## Introduction

RNA sequencing (RNA-seq) is a high throughput sequencing (HTS) method for discovery and quantification of transcripts that has become one the most employed tools in transcriptomics in recent past years (Wang et al. 2009), in biomedicine (Ergin et al. 2022), agronomy (Martin et al. 2013) or biological diversity related studies involving non-model species (López de Heredia 2016). Among the applications of RNA-seq it is worth mentioning the discovery of novel unidentified genes and splice variants (Roberts et al. 2011), the identification of functionally or adaptively relevant SNP (Zhao et al. 2019) or the assessment and quantification of the expression levels of differentially expressed genes under contrasting environmental conditions (Ekblom and Galindo 2011).

In the last years, the power of RNA-seq has grown to such extent that massive datasets are being generated for many different experimental conditions of many model and non-model species, becoming bioinformatic analyses consequently more complicated (Nazarov et al. 2017). To approach bioinformatics, many pipelines and software programs including the major steps in RNA-seq data processing (i.e. experimental design, quality control, read alignment, quantification of gene and transcript levels, visualization, differential gene expression, alternative splicing and functional analysis) have been developed. Many of these pipelines (e.g. StringTie -Pertea et al. 2016-) have been shown to be effective when working with model-organisms due to the fact that the great majority of these tools have been developed for 1000 genomes project genomes (The 1000 Genomes Project Consortium 2015). However, there is no optimal pipeline for the variety of different applications and experimental settings in which RNA-seq can be used (Conesa et al. 2016).

Although often used routinely, these available pipelines may lose their effectiveness when the species of interest has no reliable genomic reference resources available. For instance, the lack of a good quality reference genome assembly for the focal species may produce uncontrolled bias, especially if the genome assembly is highly fragmented in scaffolds or the genome includes repeated sequences, genome duplications and paralogous sequences, such as those from many conifers (Lopez de Heredia and Vázquez-Poletti 2016). In such cases, the read alignment stage will lack specificity between the query and the reference, mapping to multiple loci and introducing uncertainty in the transcript inference. In addition, the species of interest could even have no publicly available genome adding more difficulties to the RNAseq data processing. This is a common scenario when dealing with non-model species that can be partially solved by using either a *de novo* transcriptome assembly (Martin and Wang 2011) or the genome assembly of a phylogenetically close species (Raghavan et al. 2022) as a reference for read mapping and transcript quantification. *De novo* assembly is a powerful tool for species lacking a reference genome. Nevertheless, this kind of assembly relies on two implicit assumptions: (1) that the assembled transcriptome represents an unbiased, if incomplete, representation of the true underlying expressed transcriptome, and (2) that the expression estimates from the assembly are good approximations of the relative abundance of expressed transcripts. However indirect evidence suggests that these assumptions are not frequently accomplished (Freedman et al. 2020).

After the reads mapping step, transcript quantification is used for the identification of differentially expressed genes (DEGs) among distinct sample conditions, commonly known as differential expression (DE) analysis. DE is a fundamental research problem in many RNA-seq studies (Seyednasrollah et al. 2013). Several software tools based on different algorithms and normalization procedures are available to process the count matrices generated after read mapping to the reference and to conduct DE analysis. For instance, DESeq2 (Love et al. 2014) or edgeR (Robinson et al. 2009) are popular R packages that use a negative binomial distribution to model gene counts and that are integrated in many of the RNA-seq bioinformatic pipelines. Again, the selection of the proper algorithm and its corresponding parameters is of the utmost importance to get unbiased results. However, it is difficult to evaluate how modifications of the standard pipelines affect the final DE analyses results. Many benchmarking studies have been performed comparing specific software packages and methods (Seyednasrollah et al. 2015; Soneson and Delorenzi 2013; Tang et al. 2015; Baik et al. 2020; Corchete et al. 2020). However, while these studies unquestionably shed light on how the different methods work, comparing differential gene expression analysis pipeline robustness has been hampered by a lack of gold-standard datasets with known expression patterns which are required for estimating False Discovery Rates (FDR) to assess pipelines performances (Stupnikov et al. 2021).

In the present study, we report the development of the workflow DEGoldS (<u>D</u>ifferential <u>E</u>xpression analysis pipelines benchmarking workflow based on <u>Gold S</u>tandard construction). This workflow allows to test between multiple DE analysis pipelines and to select the one that produce less bias in DE inference. We further validate and show the use of this generic benchmarking workflow by applying it to standard and seven modified versions of the popular RNA-seq pipeline StringTie (Pertea et al. 2016) using as a case study real Illumina™ reads from an experiment on cambial zone tissue of radiata pine (*Pinus radiata* D. Don), a forest tree species with no publicly available reference genome yet.

## Materials and methods

### Rationale of the workflow

We designed and implemented DEGoldS, a workflow to test pipelines for DE analysis that can accommodate to a wide range of situations. DEGoldS is implemented as sequential Bash and R scripts that can run in any OS supporting UNIX, available at https://github.com/GGFHF/DEGoldS. The workflow is subdivided into four steps (Figure 1): assembly of a *de novo* transcriptome from RNA-seq reads to further exploit the produced reference transcriptome in case there is no transcriptome assembly available (Step 1); simulation of reads corresponding to transcript sets from reference transcriptome of known expression levels and to generate gold-standards (Step 2); application of the different pipelines or pipelines variations to be tested (Step 3); and assessment and selection of the best protocol by quantifying the true positives and false positives produced by each pipeline before applying it to real data (Step 4).

**Figure 1.**
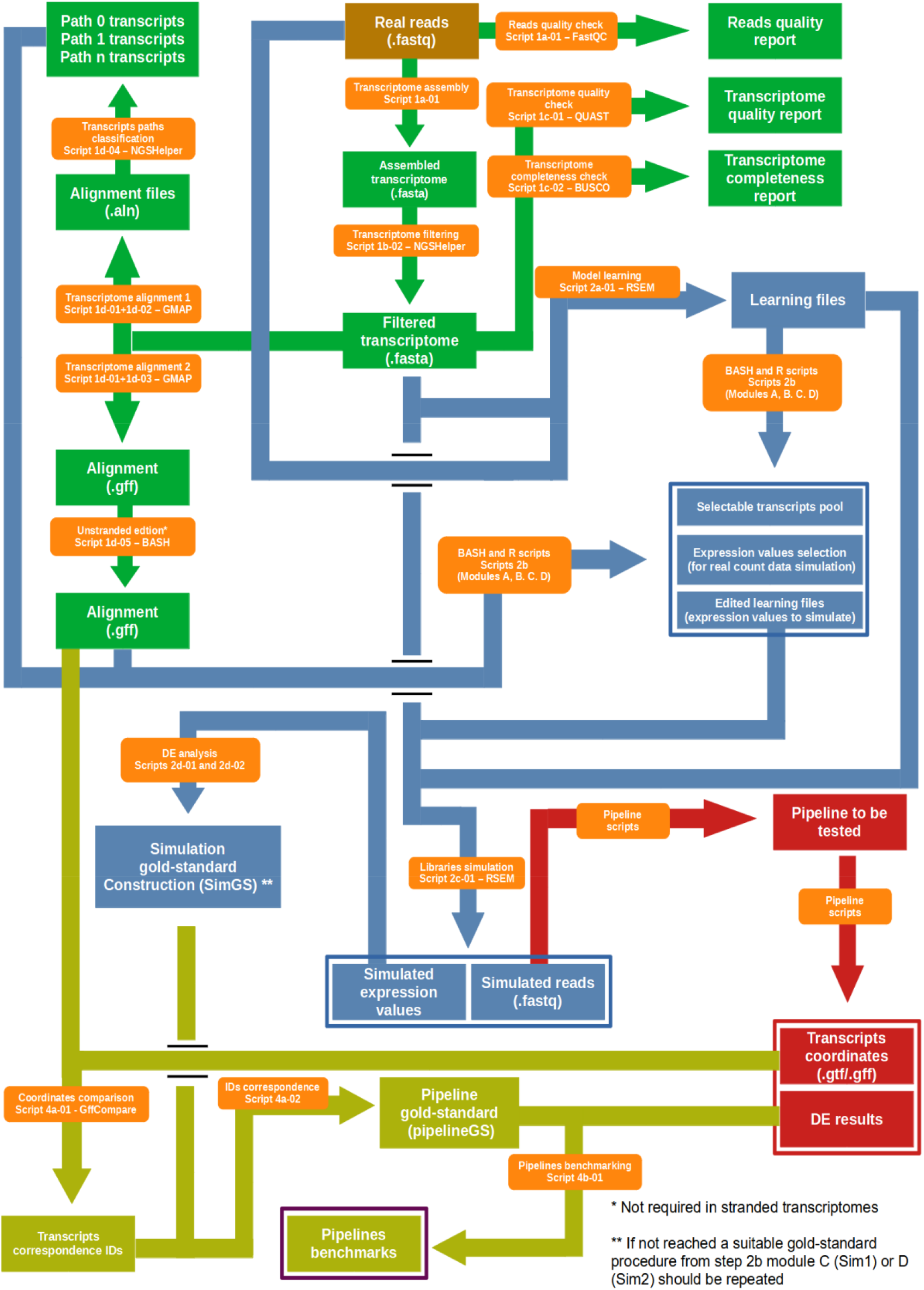
Overview of DEGolds. The processes, input and output corresponding to the four steps of the workflow are indicated: (1) transcriptome assembly (in green), (2) reads simulation (in blue), (3) tested pipelines runs (in red) and (4) pipeline benchmarking (in yellow).

**Step 1**. Reference transcriptome assembly (in case it is needed). Other available transcriptomes assemblies could be used instead. The procedure uses the same reference genome assembly from a closely related species used in the DE pipeline to be tested in order to classify transcripts of the assembled *de novo* transcriptome. The steps of this stage of the workflow are the following:

**Step 1a**. Real reads quality assessment with FastQC v0.11.9 (Andrews 2010).

**Step 1b**. *De novo* transcriptome assembly using Trinity v2.12.0 (Grabherr et al. 2011). Short and very long transcripts (<300 base pair, bp, and > 10.000 bp) are filtered out with NGShelper v0.35 (Mora-Márquez et al. 2021b).

**Step 1c**. *De novo* transcriptome quality assessment with BUSCO v4.0.6 (Manni et al. 2021) and QUAST v.5.0.2 (Mikheenko et al. 2018).

**Step 1d**. Alignment of the *de novo* reference transcriptome to the reference genome assembly of a sister species to be used in the DE pipeline of interest with GMAP (Wu and Watanabe 2005). The GMAP mapping step provides the alignment files needed for classifying transcripts into 0 times mapped (“path 0”), uniquely mapped (“path 1”) and multiple times mapped (“path n”), using NGShelper.

**Step 2**. Simulation of read files coming from selected transcripts and gold-standard generation.

**Step 2a**. Transcriptional expression profile model learning with rsem-calculate-expression function from RSEM v1.3.1 (Li and Dewey 2011). During this step, several sequencing parameters are acquired from read mapping to the reference transcriptome assembled in Step 1, including fragment length distribution, read start position distribution, sequencing error models and background noise.

**Step 2b**. Simulated transcripts and expression values selection. A user-defined number of transcripts is selected according to two categories: (1) “upregulated” transcripts that are supposed to result in DEGs between the comparison groups; and (2) “filler” transcripts that are supposed to result in non-DEGs. Both “filler” and “upregulated” transcripts are randomly selected from a pool of selectable transcripts that is delimited according to the simulation goal. In this step, the expression values in transcripts per million (TPM) that will be simulated for the “upregulated” and “filler” transcripts are also selected. RSEM outputs expression abundance information in one “.isoform.results.” file for each input library in the Step 2a. These files are edited to indicate the TPM values of interest. These TPM values are used to adjust the expression levels for each transcript in each library when simulating read files in the following step.

As explained in Step 2d, the gold-standard necessary for the benchmarking process needs to meet some requirements that are difficult to reach when working with real count data. To facilitate this process, we propose some scripts to get a selectable counts pool. This pool will then be used to select real expression profiles helping to fulfill the gold-standard requirements (see DEGoldS manual for further details).

**Step 2c**. Reads simulation. Libraries are simulated using rsem-simulate-reads function from RSEM according to selected transcripts and expression values from Step 2b and sequencing parameters from Step 2a.

**Step 2d**. Step 2c outputs “.sim.isoform.results.”, similar to “.isoform.results.” with simulated real values. These files are used to generate a gold-standard by carrying out differential gene expression analysis with the programs to be tested in Step 3. The obtained DEGs will be considered part of the gold-standard. The gold-standard needs to be composed by genes that correspond to the selected “upregulated” transcripts and should not include any gene corresponding to “filler” transcripts in order to have a robust benchmark. In case these requirements are not met, a new library simulation should be performed.

**Step 3**. The simulated read files are used as input of optimized DE pipelines to be tested with the gold-standard. In our case study, the optimized differential expression pipelines are based on the popular reference genome-guided StringTie pipeline (Pertea et al. 2016), which consists on the alignment of reads to a reference genome followed by subsequent assembly and quantification of transcripts and genes. This pipeline was subjected to modifications to optimize DE analyses in sequenced samples at three critical points (Figure 2 and Table 1) for seven combinations of modified pipelines (plus the standard pipeline). Although we are using the benchmarking workflow to test the performance of these pipelines, any other pipeline producing a GTF output could be used by adjusting the instructions in Step 3 to the alternative pipeline specificities.

**Figure 2.**
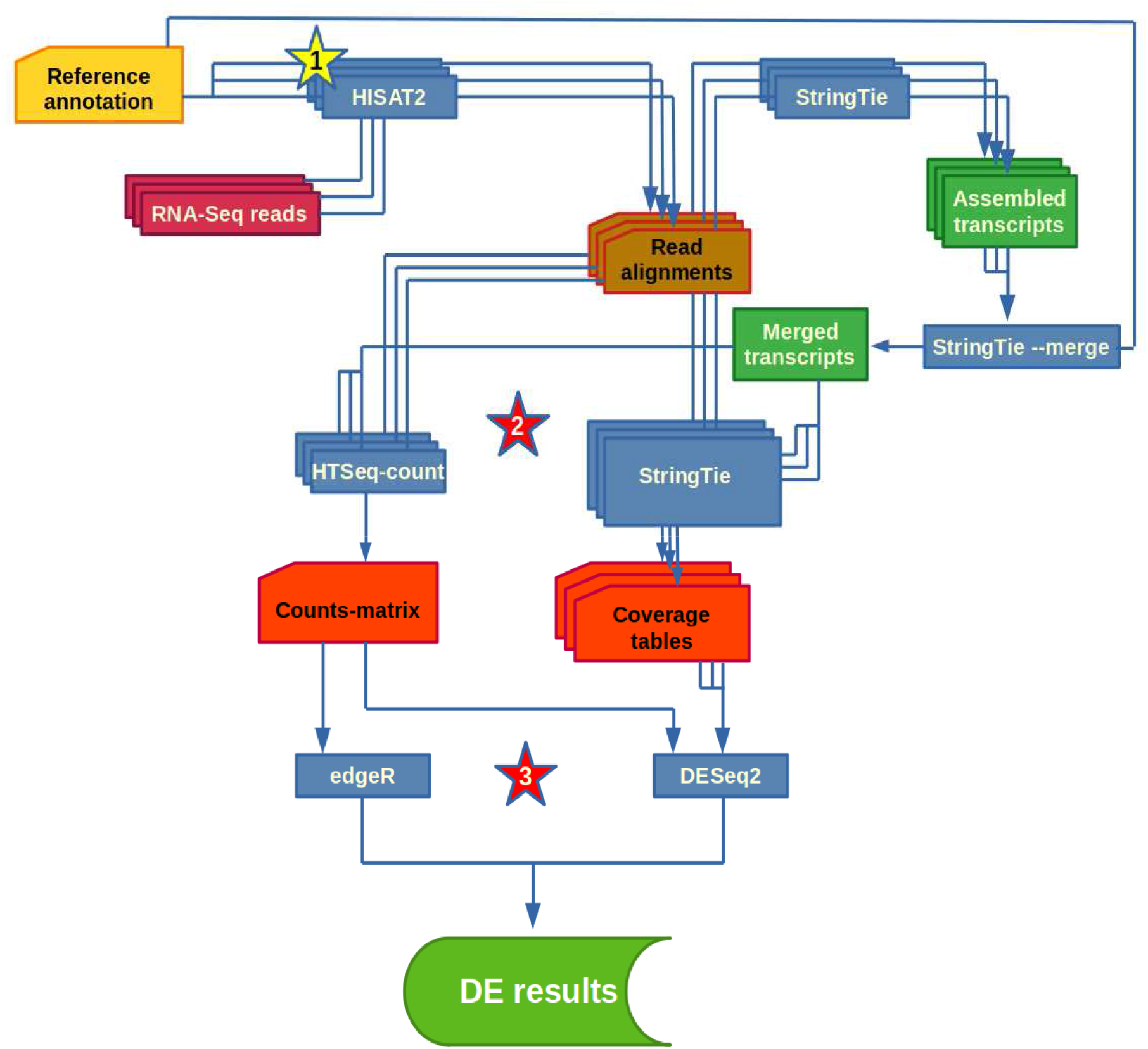
Reference-guided differential expression analyses tweaked pipeline. Image taken and modified from https://ccb.jhu.edu/software/stringtie/index.shtml?t=manual. Yellow stars represent modifications of the current StringTie pipeline in HISAT2 alignment parameters. Red stars represent additional steps to the current StringTie pipeline (HTSeq-count and edgeR).

**Table 1.** Proposed StringTie pipelines variations. Summary of the key modification to StringTie standard pipeline concerning alignment, transcript counting and DE analysis steps.

| Step | Standard pipeline | Default StringTie edgeR | Default HTSeq-count DESeq2 | Default HTSeq-count edgeR |
| --- | --- | --- | --- | --- |
| Reads alignment with HISAT2 | --score-min L,0,-0.2 (default) | --score-min L,0,-0.2 (default) | --score-min L,0,-0.2 (default) | --score-min L,0,-0.2 (default) |
| Transcript counting | StringTie | StringTie | HTseq-count | HTseq-count |
| DE analysis | DESeq2 | edgeR | DESeq2 | edgeR |
| Step | Tuned StringTie DESeq2 | Tuned Stringtie edgeR | Tuned HTSeq-count DESeq2 | Tuned HTSeq-count edgeR |
| Reads alignment with HISAT2 | --score-min L,0,-0.2 (tuned) | --score-min L,0,-0.2 (tuned) | --score-min L,0,-0.2 (tuned) | --score-min L,0,-0.2 (tuned) |
| Transcript counting | StringTie | StringTie | HTseq-count | HTseq-count |
| DE analysis | DESeq2 | edgeR | DESeq2 | edgeR |

**Step 3a**. Alignment of trimmed reads to the genome assembly of a sister species. Using HISAT2 v2.2 (Kim et al. 2019) with default parameters (except for the -dta option that was applied) can render low overall alignment rates when differences in genomic structures between the focal species and the sister species exist, or the reference genome is fragmented. For this reason, the scoring linear function parameters are tuned to increase the overall alignment rates (from the default A=-0.2 to A=-1). The resulting alignment files are processed with SAMtools v1.9 (Danecek et al. 2021) to remove duplicates and to extract concordant alignments.

**Step 3b**. Counting transcripts/genes abundance. Following the StringTie pipeline, mapped reads assembly into transcripts is carried out outputting new GTFs for each alignment (BAM) file with StringTie, using the reference GTF as a guide but allowing for novel features (not using -e option). Then, those GTF files are combined into a new “merged GTF” with the assembly of all samples. DE analyses can be performed using both input types, either transcript coverage tables or real count matrices. Coverage tables are obtained as described in the original StringTie pipeline using -eB option (t_data.ctab files). Transcript-to-gene information required in the next step for gene DE analysis is also pulled out from the same file. Count matrices are obtained from HTseq v0.12.4 package (Anders et al. 2014) using the HTSeq-count function with alignment files and “merged GTF”.

**Step 3c**. DE analysis. For gene DE analyses from coverage tables/real counts matrices two popular Bioconductor R packages are used: DESeq2 (Love et al. 2014) or edgeR (Robinson et al. 2009, McCarthy et al. 2012; Chen et al. 2016). Both DESeq2 and edgeR use a negative binomial distribution to model gene counts, but differences between them extend into several areas that could affect the final results. Coverage tables were imported into these packages with the Bioconductor R package tximport (Soneson et al. 2015).

**Step 4**. Benchmarking of the tested DE analysis pipelines.

**Step 4a**. Before the scoring a previous correspondence between Trinity and StringTie transcript IDs is needed, which is performed by trmap utility accompanying GffCompare v0.12.2 (Pertea and Pertea 2020).

**Step 4b**. Selection of the optimal pipeline: the one with more True Positives (TP) or less False Positives (FP) after comparing the DEGs obtained for the simulated read files against the gold-standard.

### Validation of the optimal pipeline selection procedure

To validate the workflow, we used a real dataset obtained from cambial zone tissue from *Pinus radiata* (https://www.ncbi.nlm.nih.gov/sra/PRJNA874614) and the public reference genome of the closely related species *P. taeda* v.2_0.1 genome assembly (Zimin et al. 2017), which is available at TreeGenes database (Falk et al. 2018; Wegrzyn et al. 2019).

#### Plant material and laboratory procedures

Cambial zone tissue samples were collected from three young (< 12 years) and three old (> 20 years) *P. radiata* trees growing in different plantations at the Basque Country (northern Spain). All trees were sampled during summer and winter periods, at noon and midnight, to maximize the number of functional genes. Young trees vascular cambial zone tissue was sampled with the aid of a leather puncher, whereas the thick bark from adult trees was peeled off with a small axe and the cambial zone tissue collected with a scalpel. Tissue samples were immediately cut into small pieces and submerged into RNA*later™* (Invitrogen) until RNA extraction.

For RNA extraction, excess RNA*later™* was blotted into paper towels and equal amounts (50 mg) of day and night samples from each individual were mixed and homogenized to a fine powder with a pestle and mortar in liquid nitrogen. Following the LiCl method (Le Provost et al. 2007) for RNA extraction, samples were immediately transferred to extraction buffer for a 10 min incubation at 65 °C. DNA was removed with DNAse I treatment and the RNA was further purified with RNA Clean & Concentrator™-25 (Zymo Research), following manufacturer’s instructions. The purified RNA was stored at -20ºC. RNA quality control, rRibosome depletion with Ribo-Zero Plant kit (Illumina), libraries construction with TruSeq Stranded Total RNA kit (Illumina) and NGS of 150 bp paired-end reads on an Illumina HiSeq 3000/4000, running HiSeq Control Software HD version 3.4.0.38, were carried out by Diagenode RNAseq services.

#### De novo P. radiata transcriptome assembly (Step 1)

Raw reads were trimmed of adapter sequences and low-quality bases (< Q30) with Cutadapt v2.10 (Martin 2011). Reads quality checks were carried out with FASTQC v0.11.9 (Andrews 2010). The 12 trimmed and quality checked libraries were employed to assemble a *de novo* reference transcriptome with Trinity v2.12.0 (Grabherr et al. 2011) using NGScloud2 (Mora-Márquez et al. 2021b) with default parameters. Short (<300 bp) and very long transcripts, (> 10.000 bp) were filtered out with NGShelper v0.535. To check for the assembled transcript quality in a *naive* way, the remaining transcripts were mapped to the *P. taeda* reference genome assembly with GMAP. The GMAP mapping step provided the alignment files needed for classifying transcripts into 0 times mapped (“path 0”), uniquely mapped (“path 1”) and multiple times mapped (“path n”), using NGSHelper. Further transcriptome quality was assessed with BUSCO v4.0.6 (Manni et al. 2021) and QUAST v.5.0.2 (Mikheenko et al. 2018). The quality of samples for DE analysis was assessed in terms of biological replicates correlation, using “Align reads and estimate abundance”, “Build expression matrix” and “RNAseq samples quality check” tools from The Galaxy platform (The Galaxy Community 2022).

#### Libraries simulation and gold-standard generation (Step 2)

In order to validate the procedure described above we used our *P. radiata* sequencing data to construct two different gold-standards for two different simulation pipelines covering from a simpler and less realistic simulation (Sim1) to a more realistic but more complex one (Sim2). Each gold-standard was set up after the differential gene expression analysis with DESeq2 and edgeR (used in the final DE analysis) with “.sim.isoform.results.” files outputted in reads simulation process. We used the *de novo* assembled transcriptome along with six winter trimmed paired-end libraries.

**Sim1**. The transcript selection step for the gold-standard consisted on the random selection of 99,900 transcripts to use as “filler” transcripts and assigned them the baseline expression level (1X=9.96 TPM). Further, 100 transcripts were randomly selected to be used as “upregulated” in the gold-standard. Half of these transcripts were upregulated in the three simulated young libraries and the other half were upregulated in the three simulated adult libraries. Expression levels of these genes were fine-tuned to ensure that all of them could be recovered as differentially expressed, while maintaining a semi-realistic situation. Expression levels were 4X, 6X, 8X, 10X and 12X, equally partitioned among the 50 transcripts. The “upregulated” transcripts were chosen from a pool of transcripts with “path 1” that did not correspond to the same *P. taeda* gene (i.e. with the same genomic coordinates). “Filler” transcripts were selected without further restrictions because they had the same TPMs for all samples so they produced no bias in gene expression due to counting isoforms within the same gene separately. Then, three libraries of 2×10^7^ paired-end reads for each group (adult *vs*. young) were simulated.

**Sim2**. Expression levels were taken from the model learning (Step 1) “.isoform.results.” files in order to get realistic expression and dispersion profiles after filtering 100 significant count values and 4,000 non significant count values for both DESeq2 and edgeR (see DEGoldS manual for further details) for accomplishing the gold-standard requirements. TPM values assigned to “upregulated” transcripts were calculated from real differentially expressed estimated count values, whereas “filler” transcripts TPM values were calculated from real non-differentially expressed estimated counts. A hundred “upregulated” and 344,647 “filler” transcripts were randomly selected from the selectable transcripts pool. In this case, all the transcripts from the “filler” pool were selected after disregarding those transcripts corresponding to the same *P. taeda* or Trinity assembly gene. The selectable “upregulated” pool was formed with the same criteria as for the “filler” pool plus the condition of being “path 1” transcripts. Finally, three libraries of 1×10^8^ paired-end reads for each group (adult *vs*. young) were simulated.

#### Genome-guided differential gene expression analysis (Step 3)

Genome-guided pipeline was applied to trimmed (real) and simulated reads together with *P. taeda* reference genome. For real reads, the initial aim in this project was to test DE analyses in all 12 samples in two factors: season (summer *vs*. winter) and age (adult *vs*. young). However, DE analyses were only conducted for the comparison between adult *vs*. young individuals using the 6 winter samples due to the very large batch effects and heterogeneity of summer samples that may bias the results. Quality check of the genome-guided alignments were carried out with HISAT2 alignment statistics as well as with results from SAMtools stats function. HISAT2 alignments were run in The Galaxy platform.

### DEGs annotation and semantic similarity

To further explore the results of DE analyses from simulated and real reads, the obtained DEGs were subjected to functional annotation followed by semantic similarity analysis. First, FASTA files with DEGs nucleotide sequences were extracted with GffRead 0.12.1 (Pertea and Pertea 2020) from the *P. taeda* reference genome with a partial “merged GTF” corresponding to the transcripts belonging to the DEGs obtained in each pipeline. Furthermore, FASTA sequences of the gold-standards were extracted from the *de novo* assembled transcriptome using NGShelper. Extracted FASTA files were annotated using TOA (Mora-Márquez et al. 2021a). TOA outputs a go-stats.csv file with all GO terms from annotated DEGs that is then used in REVIGO (Supek et al. 2011) to remove redundant GO terms. The final GO terms were used to analyze semantic similarities between pipelines with the R package GOSemSim (Yu et al. 2010, Yu 2020).

## Results

### Simulation-based workflow for DE analysis pipelines comparison

DEGoldS was implemented as a series of Bash and R scripts available at https://github.com/GGFHF/DEGoldS with the dependencies specified in them and it can run in any OS supporting UNIX. A user manual is available as a README file. The results presented below for a particular case of study showed the usability of this workflow to test any DE reference-guided pipelines, but DEGoldS can accommodate to diverse situations.

### Validation of the workflow

#### Libraries and transcriptome quality assessment

Quality checks were performed to assess de usability of the reads and the assembled *de novo* transcriptome for the bioinformatic analyses. The libraries showed differences between summer and winter samples. Firstly, the read depth of the winter libraries was about twice the summer libraries (Table S1, Suppl. File 1). Secondly, there was also an inconsistent correlation between the read depth of the biological replicates within these two groups (Figure S1, Suppl. File 1). While winter biological replicates showed uniform correlation values across all of the libraries, summer samples presented higher variation and low correlation values between them. HISAT2 alignments were also used as a read quality proxy. The overall alignment rates (OARs) were always greater for winter than for summer samples (Table S1, Suppl. File 1). After the read alignment to the *P. taeda* reference genome assembly, winter samples showed much higher coverage rates than summer ones, either with default or tuned alignment parameters (Figure S2, Suppl. File 1). Another evidence of the large variation across summer samples is the magnitude of the standard deviation in overall alignment rates (Table S1, Suppl. File 1). Therefore, the use of samples belonging only to the winter group for the differential expression analysis will avoid the hiding of DEGs, although using summer samples should not have an impact in the quality of the *de novo* transcriptome assembly, but rather increase the discovery of less represented transcripts.

The assembled *de novo* transcriptome, exhibited good quality and high transcript completeness values above 92% of complete BUSCOs (Table S2, Suppl. File 1). Then the transcriptome assembly quality was assessed by QUAST (Table S3, Suppl. File 1), with a suitable distribution of transcript lengths. Furthermore, the high number of the mapped transcripts to *P. taeda* reference genome suggested also a good quality of the *de novo* transcriptome. Indeed, only 1.28% of reads did not align to the reference genome, while 68.46% aligned once and 30.27% aligned to multiple loci.

#### Performance of DE pipelines in simulated reads

In order to analyze differences between all the modifications to the StringTie standard pipeline an *in silico* library simulation approach was adopted. After simulated reads were processed via reference-guided pipeline, much higher alignment rates were obtained when using the tuned alignment parameters in HISAT2 in contrast to the default ones in both Sim1 and Sim2 (Table 2).

**Table 2.** Simulated libraries HISAT2 OARs. AWA: Alava Winter Adult; AWY: Alava Winter Young; GWA: Guipúzcoa Winter Adult; GWY: Guipúzcoa Winter Young; VWA: Vizcaya Winter Adult and VWY: Vizcaya Winter Young.

|  | Sim1 |  | Sim2 |  |
| --- | --- | --- | --- | --- |
| Library | OAR (default) | OAR (tuned) | OAR (default) | OAR (tuned) |
| AWA | 24,77% | 45,52% | 20,02% | 42,64% |
| AWY | 23,14% | 41,36% | 19,23% | 39,12% |
| GWA | 23,79% | 43,06% | 19,38% | 40,56% |
| GWY | 23,32% | 42,11% | 19,00% | 39,75% |
| VWA | 25,71% | 44,34% | 21,69% | 41,85% |
| VWY | 22,44% | 40,50% | 18,63% | 38,23% |
| Average<br>(SD) | 23,86%<br>(1,19) | 42,81<br>(1,88) | 19,66%<br>(1,10) | 40,36%<br>(1,67) |

The total number of detected DEGs was also higher for the tuned alignment mode, especially for Sim1 and for those pipelines that included DESeq2 in Sim2 (Table 3). Overall, StringTie produced more DEGs than HTSeq-count except when edgeR was used in Sim2. For Sim2, DESeq2 showed more DEGs than edgeR for each pipeline, while the opposite pattern was found for Sim1. These differences in behavior between Sim1 and Sim2 could be explained by the unrealistic count distribution of the libraries in Sim1, as discussed below. Furthermore, most DEGs were shared between pipelines (Figure 3). Sim2 showed more exclusive DEGs for StringTie in combination with DESeq2, an expected outcome given the big amount of DEGs produced by this pipeline.

**Table 3.** Total DEGs for each performed pipeline in simulated and real libraries.

| DEGs |  | DESeq2 |  | edgeR |  |
| --- | --- | --- | --- | --- | --- |
|  |  | Stringtie | HTSeq-count | Stringtie | HTSeq-count |
| Sim1 | Default | 98 | 81 | 110 | 95 |
|  | Tuned | 175 | 143 | 200 | 163 |
| Sim2 | Default | 131 | 87 | 63 | 70 |
|  | Tuned | 410 | 140 | 99 | 109 |
| Real reads | Default | 217 | 117 | 94 | 85 |
|  | Tuned | 301 | 140 | 102 | 88 |

**Figure 3.**
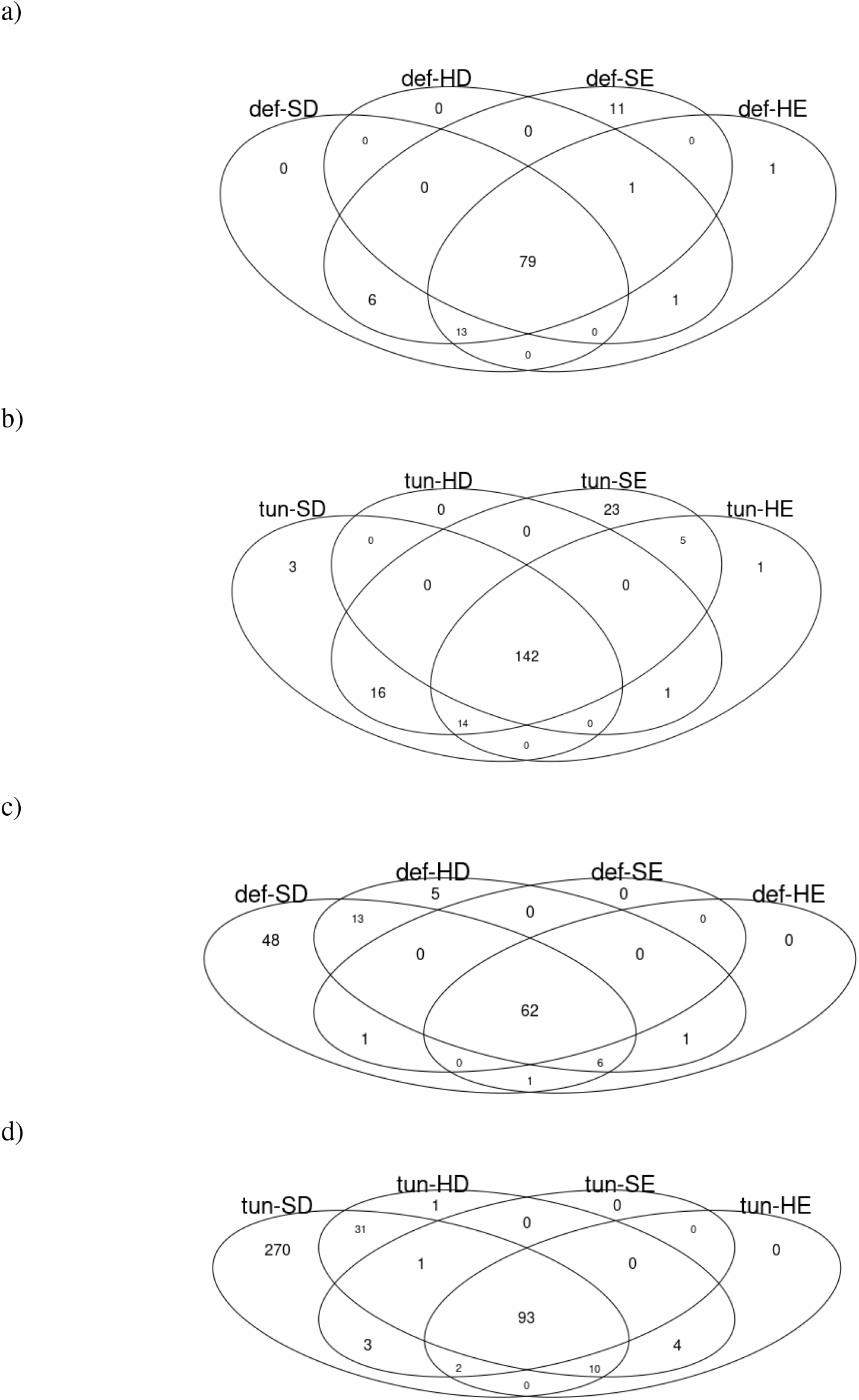
DEGs sharing in Sim1 default alignment (a), Sim 1 tuned alignment (b), Sim2 default alignment (c) and Si 2 tuned alignment between different applied reference-guided pipeline variants (def-SD=Standard pipeline; def-HD=Default HTSeq-count DESeq2; def-SE=Default StringTie edgeR; def-HE=Default HTSeq-count edgeR; tun-SD=Tuned StringTie DESeq2;tun-HD=Tuned HTSeq-count DESeq2; tun-SE=Tuned StringTie edgeR, tun-HE=Tuned StringTie edgeR)

When it comes to the scoring, the tuning of HISAT2 parameters produced a higher number of both, true positives (TP) and false positives (FP), than when using the default option (Figure 4). TPs increased around 15% for all pipelines. However the growth in FPs was massive when using StringTie (no matter DESeq2 or edgeR was used then) for Sim1 and in the combination of StringTie and DESeq2 for Sim2. When it comes to coverage/counts tables building, there were no big differences in TPs between StringTie and HTSeq-count. However, FP rate highly increased when StringTie was used for Sim1, irrespective of using DESeq2 or edgeR, and when StringTie was used together with DESeq2 for Sim2. As for the DEGs, Sim1 showed higher FP and slightly higher TP rate for edgeR than for DESeq2; however, DESeq2 had a higher number of FPs and slightly higher number of TPs for Sim2.

**Figure 4.**
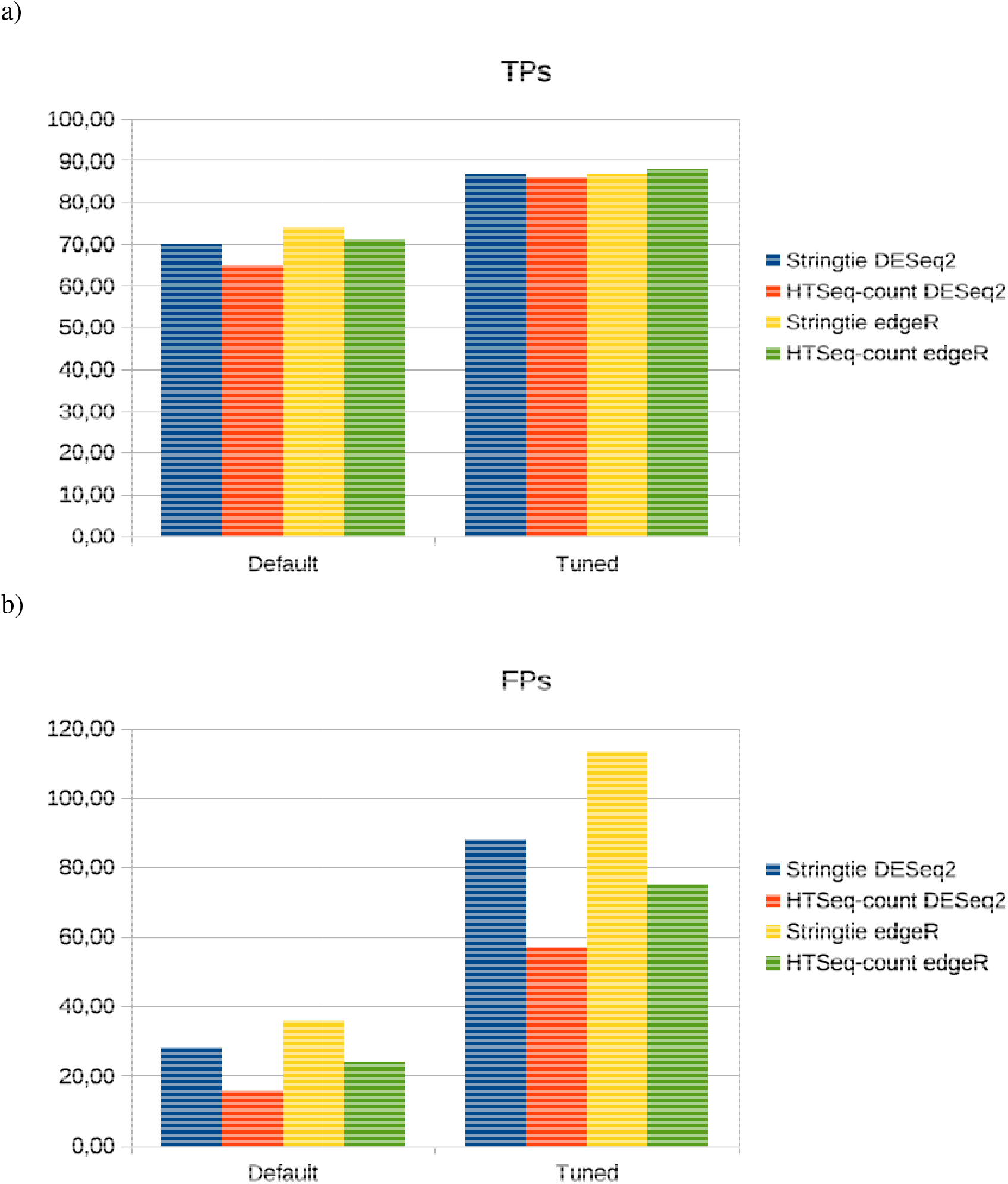
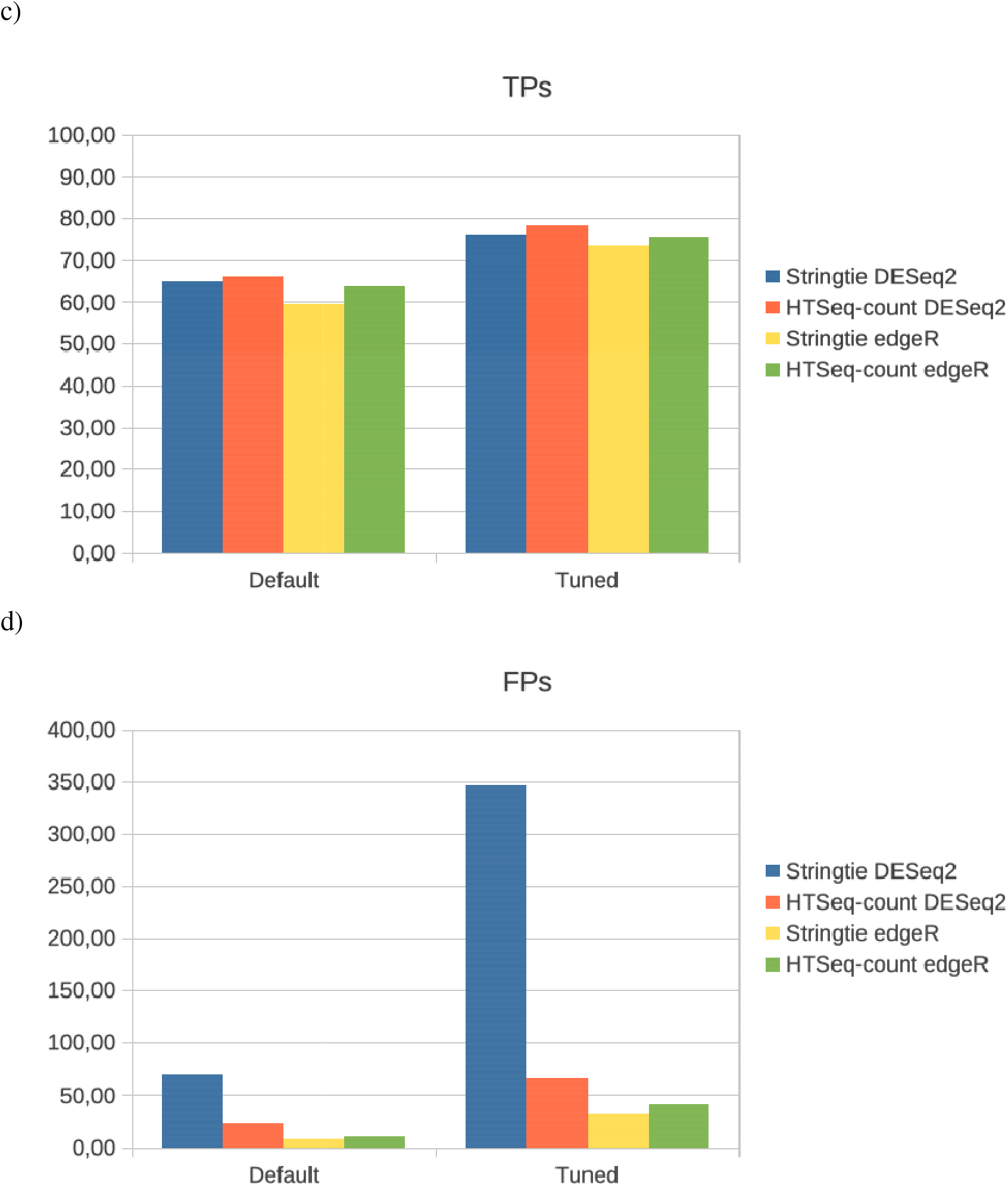
Sim1 TP (a), FP (b) and Sim2 TP (c) and FP (d) rates in genome-guided assembly differential expression results for different applied reference-guided pipeline variants.

Semantic similarity between the recovered GO terms of Biological Process (BP), Cellular Component (CC) and Molecular Function (MF) for Sim1 and Sim2 showed two clear groups between pipelines regarding to the default HISAT2 alignment mode and the tuned alignment mode (Figure 5), being the former closer to both DESeq2 and edgeR gold-standards. However no apparent differences were seen among pipeline variations within the same HISAT2 alignment mode.

**Figure 5.**
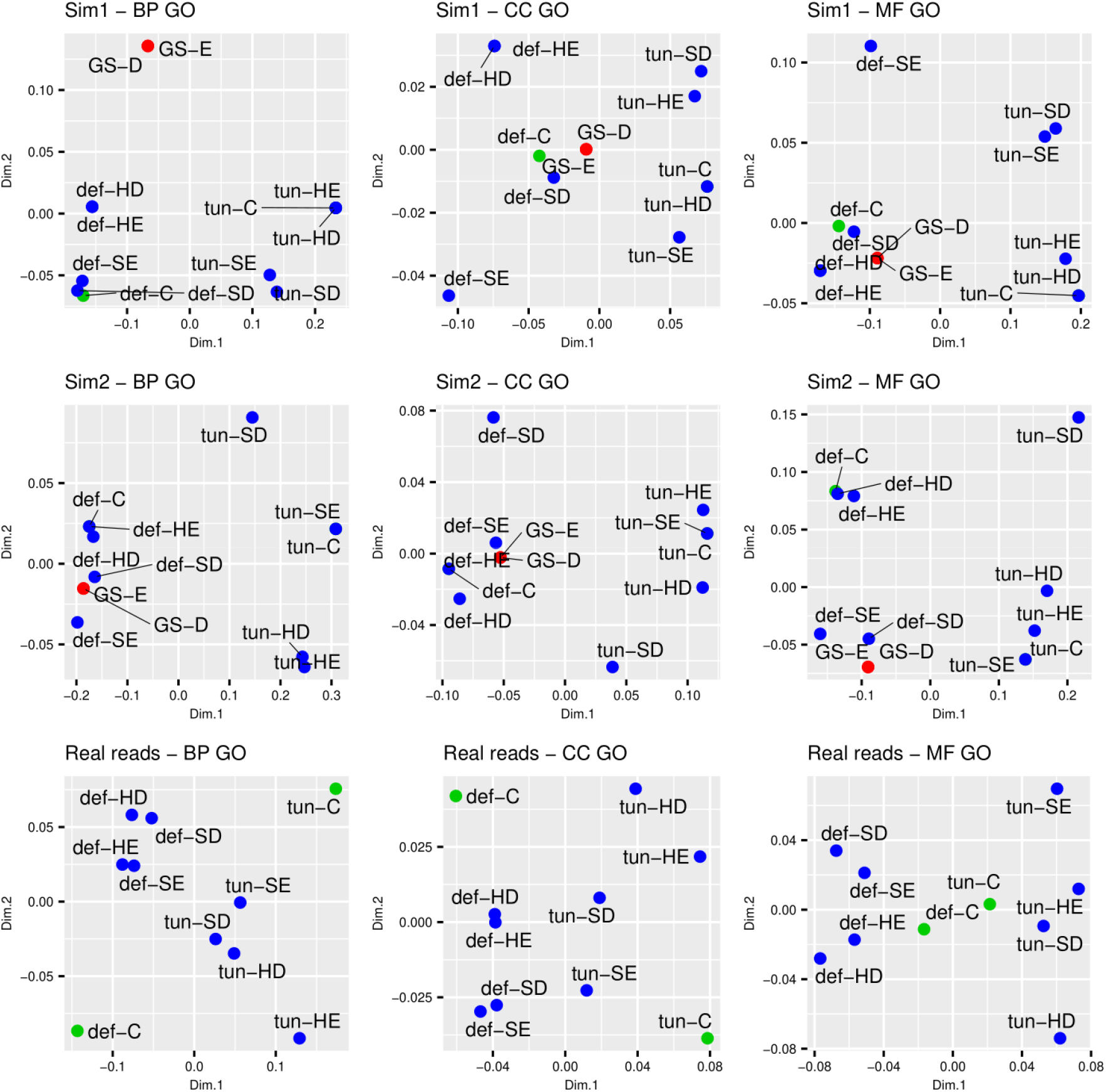
Semantic similarity between pipelines in Sim1, Sim2 and real reads for each pipeline in blue (def-SD=Standard pipeline; def-HD=Default HTSeq-count DESeq2; def-SE=Default StringTie edgeR; def-HE=Default HTSeq-count edgeR; tun-SD=Tuned StringTie DESeq2;tun-HD=Tuned HTSeq-count DESeq2; tun-SE=Tuned StringTie edgeR, tun-HE=Tuned StringTie edgeR), pipeline variations common DEGs; in green (def-C=default alignment pipeline variations; tun-C=tuned alignment pipeline variations); and gold-standards in red (GS-D=DESeq2 gold-standard; GS-E=edgeR gold-standard) for different GO terms (BP=Biological Process, CC=Cellular component and MF=Molecular Function).

#### Gold-standard generation

The benchmarking workflow could be assessed by the number of recovered DEGs in gene DE analysis using “.sim.isoforms.results” files. For Sim1 100 out of 100 of the selected “upregulated” genes were recovered using both DESeq2 or edgeR. As expected due to simulation complexity, 97 and 94 out of 100 “upregulated” selected genes were recovered for Sim2 using DESeq2 and edgeR respectively. In addition, no DEGs were reported in any cases coming from “filler” genes, which was a requirement to carry on with the benchmarking process.

#### DE pipelines with real data

Due to the high heterogeneity of the summer libraries reported before, DE analyses between young and adult trees were only performed for the six RNAseq paired-end libraries belonging to the winter group in genome-guided assembly pipelines. Consistent with the results obtained in simulations, much higher alignment rates were obtained when HISAT2 parameters were tuned. The tuned mode also recovered a larger number of DEGs than default mode, in particular when DESeq2 (Table 3). Overall, StringTie produced more DEGs than HTSeq-count. However, these differences were drastically reduced when edgeR was applied, having a similar behavior than for Sim2. Consistent again with Sim2, DESeq2 showed more DEGs, being most of them shared between pipelines, except for the StringTie-DESeq2 combination (Figure 6). Overall, the performance of the pipelines with real reads was pretty similar to Sim1 and extremely similar to Sim2. Moreover, the results for the semantic similarity between GO terms of BP, CC and MF were equal than for Sim1 and Sim2 (Figure 5): no clear differences were observed among pipeline variations within default alignment and tuned alignment modes. However, these two groups were distinguished setting up two different clusters.

**Figure 6.**
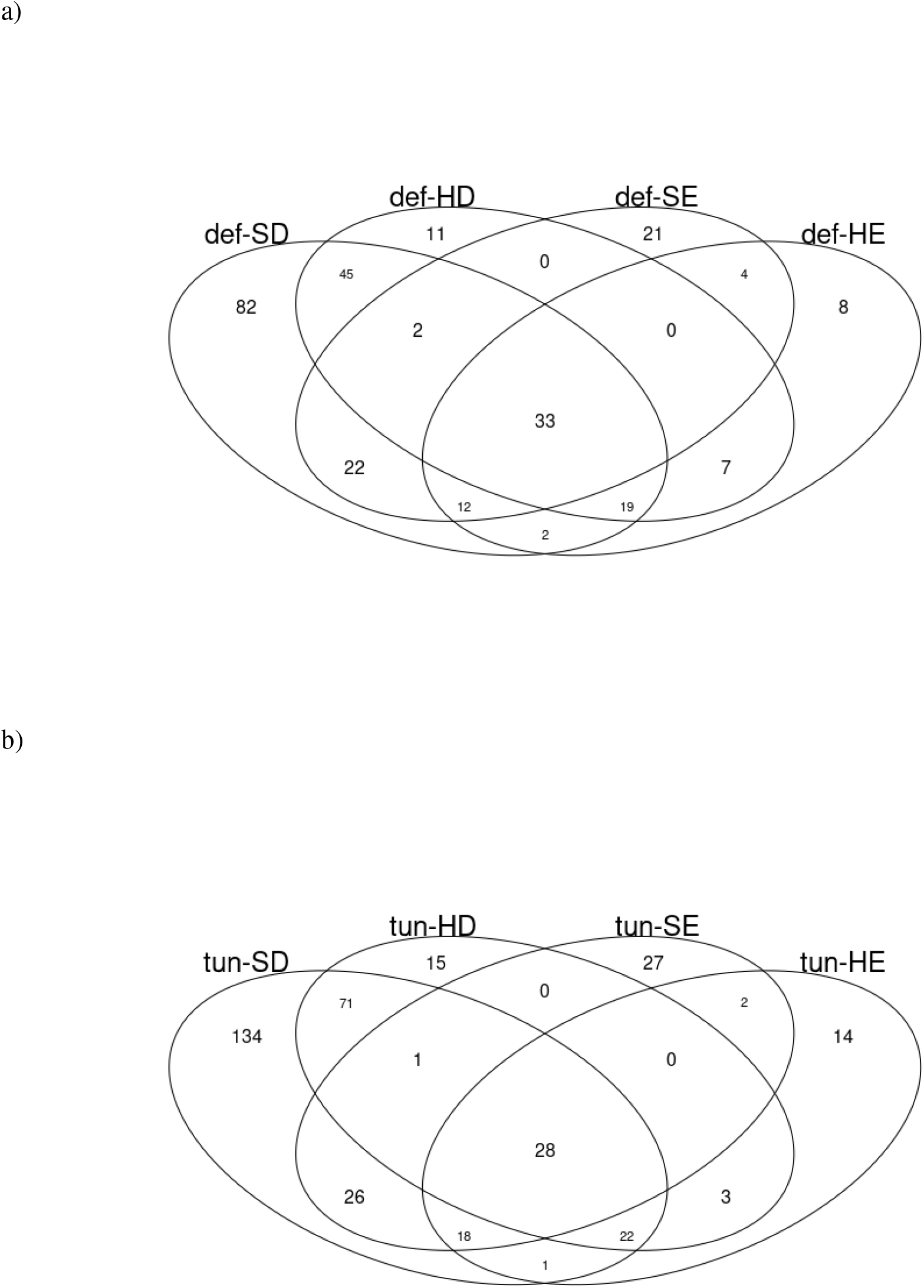
DEGs shared for real reads in the default alignment (a) and the tuned alignment (b) between different reference-guided pipeline variants.

## Discussion

The uncertainty of selecting the most appropriate pipeline and parameter settings, especially in non-model species with complex genomes that lack quality reference assemblies, is one of the biggest weaknesses of RNA-seq based DE analyses (López de Heredia and Vázquez-Poletti 2016). In general, the method returning the highest number of DEGs is considered the most suitable one because it provides more information (Liu et al. 2014). However, false positive rates should also be taken into account when looking for a strong set of results (Stupnikov et al. 2021). For this reason, there is a need to validate the applied methods as well as to understand how these alternatives to the state-of-art pipelines could affect the final results, thus helping in the search for the most suitable pipeline for each experiment. The comparison between DE analysis pipelines robustness has been hindered due to the difficulties of constructing proper gold-standard datasets with known expression values to allow the estimation of false positives (Stupnikov et al. 2021). In this study, we have presented DEGoldS, a workflow for benchmarking different DE analysis pipelines based on read simulations and a gold-standard construction. Hence, the strategy followed by DEGoldS allows to provide objective information to choose the most suitable algorithm and the best parameter combinations for the specific conditions of the RNA-seq experiment. The way RSEM utilizes the information about the expression values to simulate libraries is very suitable for the gold-standard construction (Li and Dewey 2011). The number of detected DEGs in the gold-standard was also used to assess the validity of the gold-standard and benchmarking procedure itself. The results showed a great percentage of recovered selected “upregulated” genes, even for simulations with high complexity. Moreover, those DEGs related to “filler” transcripts with low expression levels were not detected as DEGs, suggesting that this workflow is suitable for a complete DE pipeline benchmarking.

While DEGoldS can accommodate to diverse pipeline configurations, we illustrated the way it operates by testing several modifications to the widely used reference-guided StringTie pipeline (Pertea et al. 2016) and by performing two simulation scenarios: a simpler and less realistic one (Sim1) and a more realistic but more complex one (Sim2). A key difference was observed in the way DESeq2 and edgeR performed, since Sim1 exhibited more DEGs for DESeq2 than for edgeR, while for Sim2 the opposite occurred. It is important to recall that Sim1 libraries were based on a very simple TPM expression matrix, where all the transcripts had the same value for all the libraries in the “filler” transcripts, regardless the group they belonged; and the same value within the libraries of the same group in the “upregulated” transcripts. This absence of counts dispersion is used by DESeq2 and edgeR for count modeling (Love et al. 2014; Robinson et al. 2009) and, consequently, it may have an effect in the final result. To overcome this limitation, Sim2 was based on a count table obtained from the learning step of RSEM. Doing so, real count distribution and dispersion were applied, in contrast to other methods that assume a negative binomial distribution (Frazee et al. 2015). The effect of the count distribution could be probed when comparing the total DEGs obtained by DESeq2 and edgeR for real reads and for Sim2. In both cases, DESeq2 showed overall a greater number of DEGs than edgeR, conversely to Sim1. Constructing the gold-standard in Sim2 was not an easy task because trying to find DEGs from simulated transcript expression levels can lead to transcript-gene interference resulting in an unreliable gold-standard. However, the implemented scripts in DEGolds overcome this limitation, enabling the construction of a robust gold-standard even with real count data. Actually, Sim2 gathers up more complexities from experimental data thus simulating more realistically the particular conditions of real experiments.

When it comes to StringTie pipeline benchmarking, large differences on the DE outcome were observed between all the variations applied to the standard conditions, suggesting that any single variation of the pipeline could have a large effect on the final results of DE, both quantitatively (number of DEGs) and qualitatively (distribution of the recovered GO terms, TP and FP). Interestingly, the alignment step revealed to be critical to increase the TPs but at the expense of increasing also the FPs. Furthermore, there was also an impact of the parameter tuning of HISAT2 on the distribution of the recovered GO terms, as revealed by the two big clusters obtained in the semantic similarity multi-dimensional scaling (Figure 5) that suggested a better performance using the default parameters than the tuned ones. These results suggest that the parameter tuning is not necessarily the best choice, even when the reference genome is highly fragmented and does not correspond to the focal species. Therefore, taking all the simulation results into account, we can conclude that the tuning alignment step should only be considered if the final objective of the study is to find more TP DEGs, no matter a collateral increase of FPs.

Beyond the alignment step, it was observed that the identified DEGs were mostly shared by all the pipelines, except for the cases were the total number of DEGs increased significantly, and that TPs hardly varied between pipelines. This means that the impact of modifying the standard pipelines was nearly exclusive in terms of FPs, suggesting that recovering less DEGs is more appropriate than recovering large numbers of DEGs that can be spurious, in contrast to some studies that claim that the more DEGs the better (Liu et al. 2014). Actually, while the original pipeline proposes the use of StringTie together with DESeq2, this combination showed to be the worst comparing to the other pipeline modifications in terms of FP. Conversely, the FP ratio decreased noticeably when HTSeq-count and DESeq2 were used together or when edgeR was used in combination either with StringTie or HTSeq-count.

## Conclusions

In this study we propose a simulation-based workflow for DE analysis pipeline benchmarking (DEGoldS) that simulates a realistic gold-standard using expression values learned from real data, allowing objective tests of the performance of different user-defined DE pipeline modifications in terms of True and False Positives. We applied DEGoldS to a popular StringTie reference-guided DE pipeline to illustrate the validity of the workflow and to show how modifications of a standard pipeline can significantly vary the expected results, thus increasing the quality and reliability of the results. Moreover, we have shown that less DEGs could mean more robust results, contrary to the usual thought of more DEGs meaning better results.

## Acknowledgments

This study has been carried out in memoriam of Dr. Pablo G. Goicoechea, who initiated this research and participated in the initial design of the work.

