## Supplemental File 1 for "DEGoldS: a workflow to assess the accuracy of differential expression analysis pipelines through gold-standard construction"

**Table S1, Suppl. File 1.** Real libraries read depth after trimming and their HISAT2 OARs. ASA: Alava Summer Adult; ASY: Alava Summer Young; GSA: Guipúzcoa Summer Adult; GSY: Guipúzcoa Summer Young; VSA: Vizcaya Summer Adult and VSY: Vizcaya Summer Young. AWA: Alava Winter Adult; AWY: Alava Winter Young; GWA: Guipúzcoa Winter Adult; GWY: Guipúzcoa Winter Young; VWA: Vizcaya Winter Adult and VWY: Vizcaya Winter Young.

| Library | Number of read pairs | OAR (default) | OAR (tuned) |
| --- | --- | --- | --- |
| ASA | 73541086 | 6,43 % | 60,25 % |
| ASY | 82352648 | 7,48 % | 66,84 % |
| GSA | 80998614 | 8,5 % | 60,96 % |
| GSY | 71651324 | 30,97 % | 70,35 % |
| VSA | 72854930 | 25,63 % | 68,73 % |
| VSY | 74373616 | 6,97 % | 60,54 % |
| AWA | 113493192 | 68,73 % | 87,89 % |
| AWY | 101347482 | 74,48 % | 92,63 % |
| GWA | 88822392 | 71,61 % | 91,52 % |
| GWY | 106970166 | 71,76 % | 91,97 % |
| VWA | 126182445 | 30,44 % | 71,15 % |
| VWY | 101603001 | 68,67 % | 89,75 % |
| All  (SD) |  | 39,31 %  (29,37 %) | 76,05 %  (13,51 %) |
| Summer  (SD) |  | 25,13 %  (25,09 %) | 69,85 %  (11,58 %) |
| Winter  (SD) |  | 64,28 %  (16,72 %) | 87,48 %  (8,19 %) |

**Table S2, Suppl. File 1.** Transcriptome completeness assessment with BUSCO statistics.

|  | Total | % |
| --- | --- | --- |
| Complete BUSCOs | 1491 | 92,40 % |
| Complete and single-copy BUSCOs | 476 | 29,50 % |
| Complete and duplicated BUSCOs | 1015 | 62,90 % |
| Fragmented BUSCOs | 28 | 1,70 % |
| Missing BUSCOs | 95 | 5,90 % |
| Total BUSCO groups searched | 1614 |  |

**Table S3, Suppl. File 1.** Transcriptome assembly quality assessment with QUAST statistics.

| # contigs (>= 0 bp) | 1637886 |
| --- | --- |
| # contigs (>= 1000 bp) | 243968 |
| # contigs (>= 5000 bp) | 13471 |
| # contigs (>= 10000 bp) | 1 |
| # contigs (>= 25000 bp) | 0 |
| # contigs (>= 50000 bp) | 0 |
| Total length (>= 0 bp) | 1178253686 |
| Total length (>= 1000 bp) | 542420676 |
| Total length (>= 5000 bp) | 86233498 |
| Total length (>= 10000 bp) | 10000 |
| Total length (>= 25000 bp) | 0 |
| Total length (>= 50000 bp) | 0 |
| # contigs | 630389 |
| Largest contig | 10000 |
| Total length | 800989753 |
| GC (%) | 38.81 |
| N50 | 1632 |
| N75 | 826 |
| L50 | 131713 |
| L75 | 308341 |
| # N's per 100 kbp | 0,00 |

**a)**

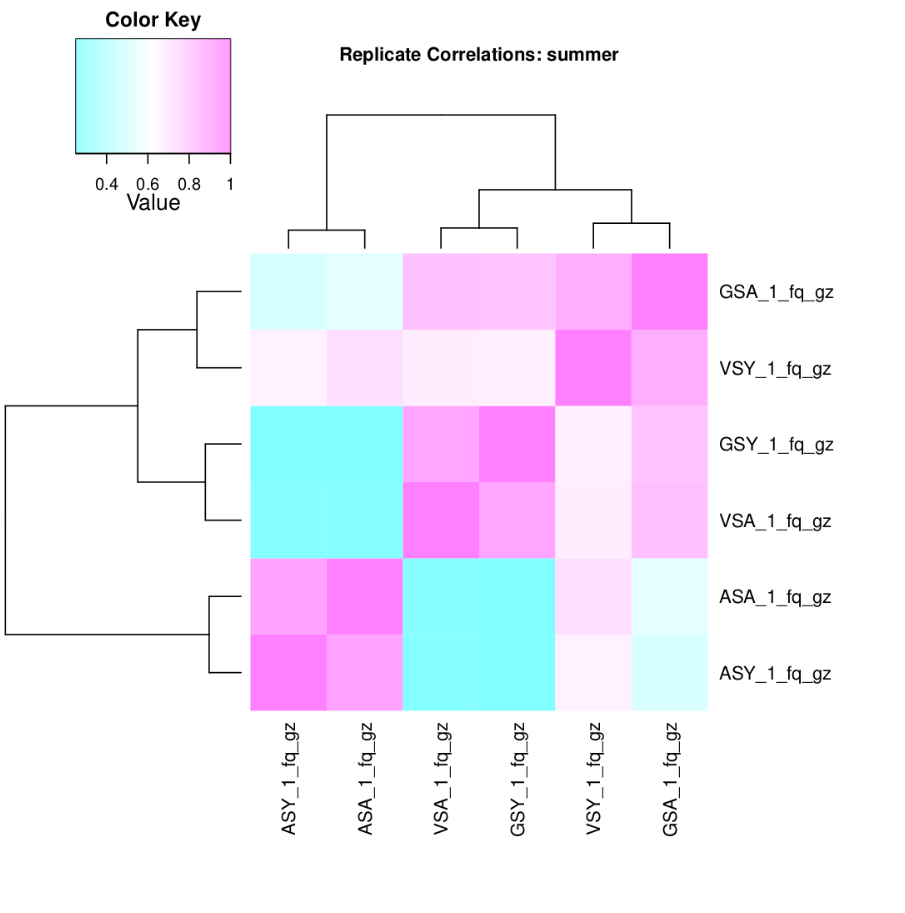

**b)**

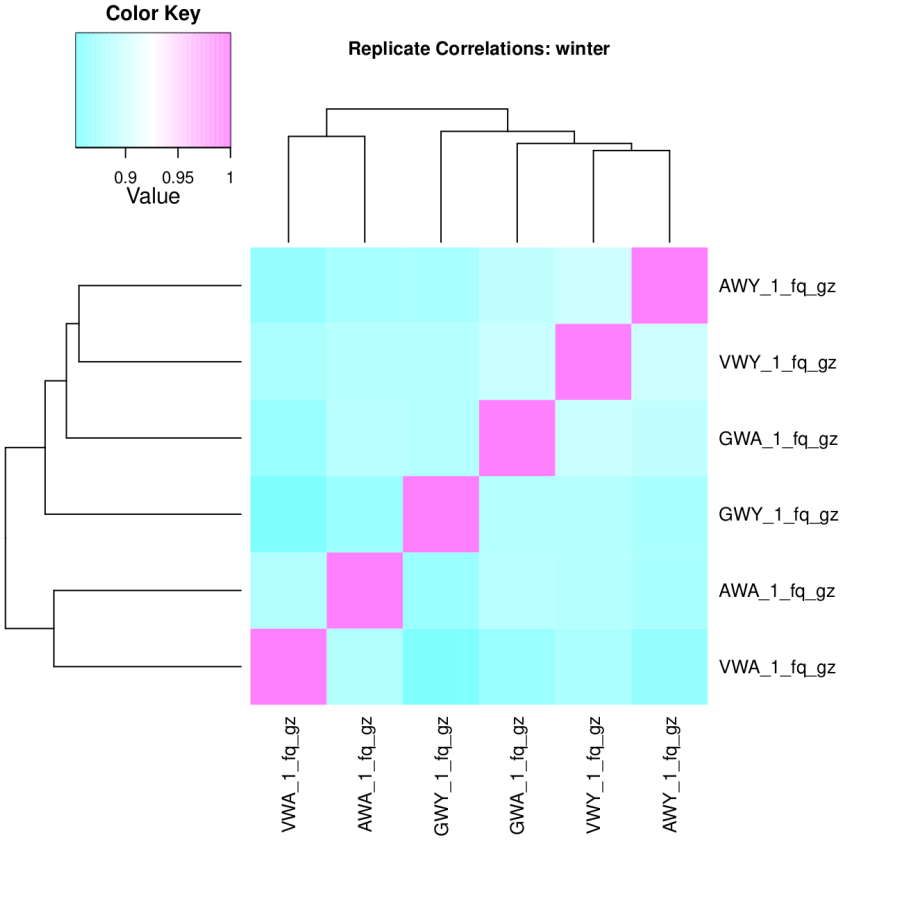

**Figure S1, Suppl. File 1.** Summer samples show a very low correlation between biological replicates (a) which could led to DEGs hiding. On the contrary winter samples have a very high correlation values allowing the differential expression analysis based on these 6 samples. ASA: Alava Summer Adult; ASY: Alava Summer Young; GSA: Guipúzcoa Summer Adult; GSY: Guipúzcoa Summer Young; VSA: Vizcaya Summer Adult and VSY: Vizcaya Summer Young. AWA: Alava Winter Adult; AWY: Alava Winter Young; GWA: Guipúzcoa Winter Adult; GWY: Guipúzcoa Winter Young; VWA: Vizcaya Winter Adult and VWY: Vizcaya Winter Young.

a)
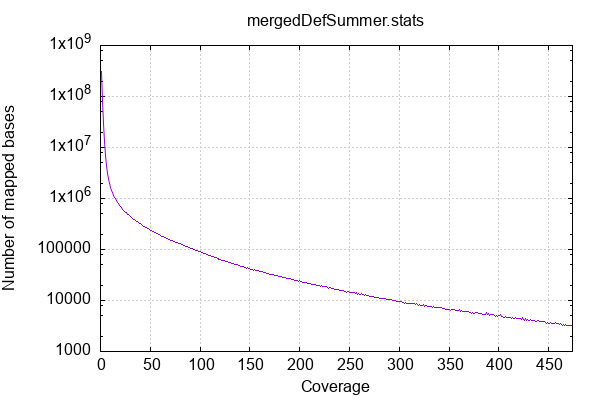
b)
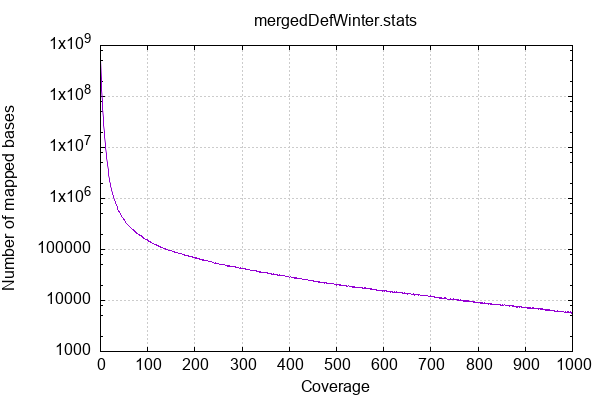

c)
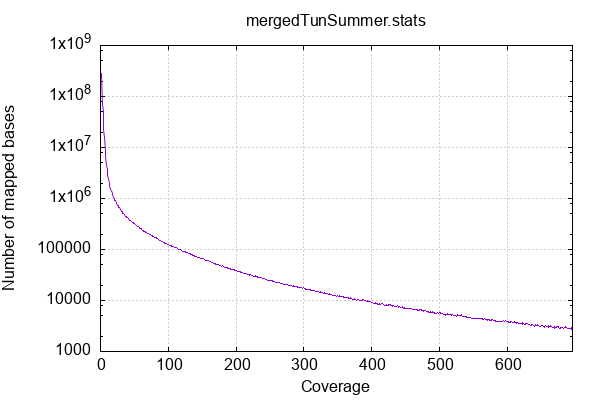

d)

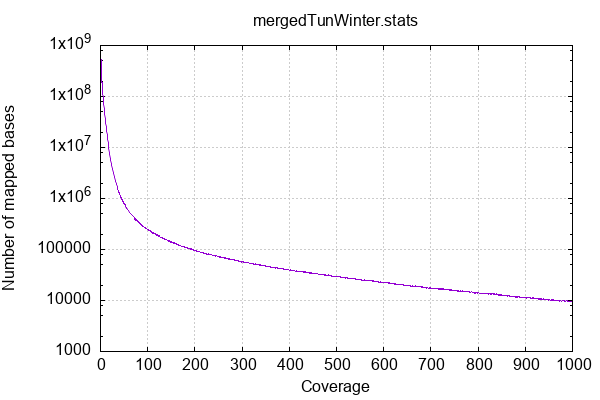

**Figure S2, Suppl. File 1.** Summer samples merged default alignment (a) has lower coverage values than winter samples merged tuned alignment (c). In the same way, summer samples merged tuned alignment coverage (b) is lower than winter samples merged tuned alignment (d)
